# A Straightforward and Robust Enzymatic Reporter System for Anaerobic Thermophiles

**DOI:** 10.1101/2025.06.23.661153

**Authors:** Joey L. Galindo, Hansen Tjo, Jonathan M. Conway

## Abstract

Thermophilic anaerobic organisms, particularly species that can naturally degrade lignocellulosic biomass, show great promise for next generation bioprocessing. This has led to the development of nascent genetic systems to metabolically engineer these non-model organisms. However, a major challenge remains a lack of reliable reporter systems compatible with the combination of thermophilic and anaerobic growth conditions. Additionally, native glycoside hydrolases in these organisms limit the usefulness of traditional glycosidic enzyme reporters (e.g. LacZ) because of the native background activity present on para-nitrophenyl glucoside substrates. Here we describe the development of a straightforward and robust enzymatic reporter system that overcomes these challenges in *Anaerocellum* (f. *Caldicellulosiruptor*) *bescii,* an anaerobic, extremely thermophilic (T_opt_ ∼78 °C), lignocellulolytic bacterium. Our method is based on heterologous expression of hyperthermophilic archaeal galactosidases: an α-galactosidase from *Pyroccous furiosus* (*Pf*αgal), and a β-galactosidase from *Caldivirga maquilingensis* (*Cm*βgal). We show that these reporters produce strong, orthogonal signals on colorimetric substrates at high temperatures (≥90°C) that eliminate background activity from endogenous galactosidases. We then demonstrate the capability of *Cm*βgal, the stronger of the two reporters, to distinguish differences in levels of expression between *A. bescii* promoter sequences, which we verify through qRT-PCR. With its high signal to noise ratio and ease of use, this reporter system offers a reliable method for assessing protein expression in anaerobic thermophilic organisms, opening doors to improved genetic tools and metabolic engineering applications for industrial biotechnology.

## Introduction

Reducing the world’s dependence on non-renewable and geographically limited fossil fuel-based feedstocks is a critical challenge. One promising alternative feedstock is plant biomass, especially its most common form, lignocellulose, which could provide an inexpensive and plentiful source of renewable energy and industrial chemicals (Lynd *et al*., 2022; Langholtz *et al*., 2024). The recalcitrance of lignocellulosic biomass severely limits its utilization through conventional bioprocessing approaches (Bing *et al*., 2021; Lynd *et al*., 2022). However, several thermophilic anaerobic bacteria are capable of natively breaking down lignocellulose, making them prime candidates for metabolic engineering (Blumer-Schuette *et al*., 2014; Lee *et al*., 2020; Bing *et al*., 2021). Yet, the genetic toolkits available in these non-model bacteria are still extremely limited, which has hampered engineering efforts (Loder *et al*., 2017; Blumer-Schuette, 2020). A major roadblock impeding the development of genetic tools in these thermophilic anaerobic bacteria is the lack of easily observable and background-free reporter systems that are compatible with the high temperature and oxygen-free growth conditions of these organisms (Loder *et al*., 2017; Riley and Guss, 2021; Streett, Charubin and Papoutsakis, 2021).

*Anaerocellum* (f. *Caldicellulosiruptor*) *bescii* is the most thermophilic lignocellulose degrading bacteria known, with an optimal growth temperature of 75-78 °C under anaerobic conditions (Lee *et al*., 2020). Development of genetic tools in this organism have enabled the metabolic engineering of *A. bescii*. These tools include deletions in the *pyr* locus (either Δ*pyrF* or Δ*pyrE*) to create uracil auxotroph strains that allows for positive selection with *pyr* gene complementation and counter selection on 5-FOA for marker replacement in *A. bescii* (Cha *et al*., 2013; Lipscomb *et al*., 2016). Positive selection is also available using a highly thermostable kanamycin resistance gene (*htk*) and selection on kanamycin antibiotic (Lipscomb *et al*., 2016).

Using these tools, *A. bescii* has been successfully engineered to produce several industrially relevant products including ethanol, acetone, and 2,3-butanediol (Bing *et al*., 2024; Straub *et al*., 2020; Tanwee *et al*., 2023). However, the lack of robust, well-characterized genetic parts (e.g., promoters, reporters, terminators) as part of this genetic toolkit in *A. bescii* remains a major limitation to expanding metabolic engineering in it and similar organisms.

Control over protein expression is often most effectively achieved at the transcriptional level by varying the specific promoter sequence upstream of a gene to change the level transcribed by RNA polymerase (Kim, Sinnott and Sandoval, 2020; Riley and Guss, 2021). Yet, to date, expression of heterologous proteins in *A. bescii* has relied almost exclusively upon three native constitutive promoters associated with genes for the S-layer protein (P_slp_), a S30 ribosomal protein (P_S30_), and a bifurcating-hydrogenase (P_bh_) (Lee *et al*., 2020; Tanwee *et al*., 2023, 2023; Bing *et al*., 2024). All of these promoters are thought to drive relatively high expression, but there have been no direct comparisons of their strengths at the protein level (Lipscomb *et al*., 2016; Williams-Rhaesa *et al*., 2018; Lee *et al*., 2020). Furthermore, other methods of modulating transcription like CRISPRi, which has been demonstrated in other thermophiles, have yet to be implemented in *A. bescii* (Ganguly, Martin-Pascual and van Kranenburg, 2020; Riley and Guss, 2021). A suitable anaerobic, extremely thermophilic protein reporter system would greatly enhance efforts to develop these and other genetic engineering tools in *A. bescii*.

Finding protein-based reporters that work well in anaerobic thermophiles has proven challenging because many reporter proteins permanently denature at the high native growth temperatures of thermophilic bacteria (Jensen *et al*., 2017; Kim, Sinnott and Sandoval, 2020; Riley and Guss, 2021; Hocq *et al*., 2023). Furthermore, many fluorescent or luminescent reporter proteins, like GFP and luciferase, require oxygen to activate, and thus cannot be used in strict anaerobic conditions (Kim, Sinnott and Sandoval, 2020; Riley and Guss, 2021; Streett, Charubin and Papoutsakis, 2021; Hocq *et al*., 2023). Other fluorescent proteins like flavin mononucleotide (FMN)-binding fluorescent proteins (FbFPs), can fluoresce anaerobically under blue light but are quite dim compared to conventional fluorescent reporters (Kim, Sinnott and Sandoval, 2020; Riley and Guss, 2021; Streett, Charubin and Papoutsakis, 2021). Another option is a class of protein tags which produce light upon binding to a small molecule ligand, the most notable of which are Snap-Tag, Clip-Tag, Halo-Tag, and Fluorescence-Activating Absorption-Shifting Tag (FAST) (Kim, Sinnott and Sandoval, 2020; Riley and Guss, 2021; Streett, Charubin and Papoutsakis, 2021; Hocq *et al*., 2023). However, most of these tags are not thermostable enough to be used in extreme thermophiles like *A. bescii,* or have shown limited practical use *in vivo* (Mattossovich *et al*., 2020; Merlo *et al*., 2022; Hocq *et al*., 2023). The most promising demonstration of these fluorescent protein tags was by Hocq *et al*. who show certain FAST tag variants had melting temperatures of ∼70 °C *in vitro* (Hocq *et al*., 2023). But, when expressed in an anaerobic thermophilic bacterium, *Thermoanaerobacter kivui*, the reporter only functioned effectively *in vivo* up to 55 °C (Hocq *et al*., 2023). Furthermore, these tags require the addition of fluorogenic dyes for detection, some of which require added wash steps, which complicate reporter readout (Riley and Guss, 2021; Streett, Charubin and Papoutsakis, 2021; Hocq *et al*., 2023).

An alternative to fluorescent proteins is enzymatic reporters, such as the widely used *E.coli* β-galactosidase (*lacZ*) and β-glucuronidase (*gusA*) based systems, which detect protein expression indirectly by breaking down precursor molecules to produce a quantifiable change in colored product (Kim, Sinnott and Sandoval, 2020; Riley and Guss, 2021; Streett, Charubin and Papoutsakis, 2021). These systems have been used extensively in mesophilic anaerobes since many of these colorimetric molecules, like various ortho- or para-nitrophenol linked compounds, do not require oxygen to produce a change in color (Jensen *et al*., 2017; Streett, Charubin and Papoutsakis, 2021). A number of thermostable versions of these enzymes have been identified, but their implementation as reporters has remained limited (Honarbakhsh *et al*., 2012; Fujita *et al*., 2015; Jensen *et al*., 2017; Loder *et al*., 2017). This is in part because many thermophilic bacteria, particularly species that possess large inventories of lignocellulolytic enzymes, often express native versions of these enzymatic reporters or enzymes with identical activity, resulting in background activity that obscures any signal from the reporter (Honarbakhsh *et al*., 2012; Fujita *et al*., 2015). Thus, most attempts to implement enzymatic reporter systems in thermophiles have required time consuming deletions of the native enzyme from the genome or heterologous expression in species that do not produce background activity (Honarbakhsh *et al*., 2012; Fujita *et al*., 2015; Jensen *et al*., 2017; Loder *et al*., 2017). Some notable attempts to implement enzymatic reporters in extreme thermophiles include expression of a β-glucuronidase in the archaeon *Sulfolobus solfataricus,* a β-galactosidase in the bacterium *Thermus thermophilus,* and a β-galactosidase from *Geobacillus stearothermophilus* in *Geobacillus thermoglucosidasius*, none of which are obligate anaerobes (Honarbakhsh *et al*., 2012; Fujita *et al*., 2015; Jensen *et al*., 2017). Thus far, the most practical attempt to implement a thermostable enzymatic reporter in an anerobic thermophile was by Olsen *et. al*, who used the aforementioned β-galactosidase from *G. stearothermophilus* to characterize promoters in the anaerobic moderate thermophile *Acetivibrio thermocellus* (fm. *Clostridium thermocellum*), which like *A. bescii* is also highly efficient at degrading lignocellulose (Olson *et al*., 2015). However, compared with our method this required a more involved preparation of cell extracts, and did not explicitly eliminate background activity from endogenous galactosidases.

Here, we demonstrate a new reporter system for lignocellulolytic thermophiles using hyperthermophilic galactosidases: an α-galactosidase from *Pyroccous furiosus* (*Pf*αgal, T_opt_ =115°C), and a β-galactosidase from *Caldivirga maquilingensis* (*Cm*βgal, T_opt_=110°C) (van Lieshout *et al*., 2003; Letsididi *et al*., 2017). The optimal temperatures of these reporter enzymes are above the temperature where native *A. bescii* enzymes are stable, thus enabling the elimination of background activity with a ≥90°C incubation. The resulting reporter assay, consisting of a heat inactivation step followed by incubation with pNP-galactopyranoside substrate, produces a strong colorimetric signal while eliminating background from native enzymes. We demonstrate the utility of this reporter system by using it to compare the protein expression driven by two previously utilized *A. bescii* promoters. We validate that these protein expression results align with the transcriptional levels driven by these promoters. Together, this system offers a powerful reporter tool for the analysis of genetic parts and genetic manipulations in *A. bescii.* Furthermore, these reporters could easily be adapted for use in other anaerobic thermophiles of interest as microbial chassis for industrial biotechnology.

## Materials and Methods

### Bacterial strains and growth conditions

Plasmids were cloned in chemically competent *Escherichia coli* 10beta (New England Biolabs) or TOP10 (Thermo Scientific). *E. coli* cultures were maintained at 37 °C in enriched Luria-Bertani (LB) medium (24 g/L yeast extract, 10 g/L tryptone, 5 g/L NaCl) or LB agar medium (5 g/L yeast extract, 10 g/L tryptone, 5 g/L NaCl, 15 g/L agar) plates with 50 µg/ml apramycin (Thermo Scientific). Unless described as otherwise, *A. bescii* strains were cultured in 50 mL of medium in 125 ml serum bottles sealed with 20 mm butyl rubber stoppers at 70 °C without shaking. These strains were grown in modified complex DSM 516 media containing 0.5 g/L yeast extract and 5 g/L maltose substrate, referred to following convention as CM516 medium, and supplemented with 50 µg/ml kanamycin (IBI Scientific) as appropriate, referred to as CM516K medium (Lipscomb *et al*., 2016). Sealed serum bottles containing sterile medium were made anaerobic through vacuum and gas cycling, with the headspace being replaced with 80% (v/v) N_2_ and 20% (v/v) CO_2_ gas. As is standard, *A. bescii* cell density was measured as the optical density at 680 nm (OD680) using a cuvette in a Nanodrop One C spectrophotometer (Thermo Scientific) with 1x DSM 516 salt solution used as the blanking solution (Lipscomb *et al*., 2016; Rodionov *et al*., 2021; Tjo *et al*., 2025).

### Vector construction

Tables of oligonucleotide primers and synthesized DNA used to construct the plasmids in this study can be found in the supporting information (**Table S1&2**). The two promoter sequences used to express the reporter genes in this study consisted of the 200 bp sequences immediately upstream of the start codon of their associated gene (**Table 1**). These promoters were P_slp_ associated with the S-layer protein gene (*Athe_2303*), and P_bh_ associated with a bifurcating-hydrogenase gene (*Athe_1295*) (**Table 1**). Maps of plasmids constructed and utilized in this study are shown in **Figure 1**. pSBS4 (Empty Vector) was obtained from the Dr. Robert Kelly lab (North Carolina State University) (Lipscomb *et al*., 2016). This vector consists of a native *A. bescii* replicating plasmid (pAthe02), the *htk* gene expressed by promoter P_S30_ associated a S30 ribosomal protein (Athe_2105), as well as elements that enable cloning in *E. coli* including: an apramycin resistance marker (Apr), replication initiation protein A (repA), and the pSC101 origin (**Fig. 1**) (Chung *et al*., 2013; Lipscomb *et al*., 2016). Vectors pJLG091 and pJLG093 express the α-galactosidase from *Pyroccous furiosus* (*Pf*αgal) and the β-galactosidase from *Caldivirga maquilingensis* (*Cm*βgal) respectively with P_slp_ (**Fig. 1**). This expression site is based on the protein expression construct used previously in pJMC046 with the P_slp_ promoter and Calkro_0402 terminator, but is relocated on the pSBS4 backbone between Apr and pAthe02 (Conway *et al*., 2018). The backbone DNA for these vectors was PCR amplified from a sequenced plasmid that had been constructed previously via the insertion of a different P_slp_ driven gene into the pSBS4 backbone at this same site (**Table S1;** Primers JLG021-22). Codon optimized genes flanked by appropriate overlapping regions were purchased (Twist Biosciences) for *Pf*αgal and *Cm*βgal (**Table S2**) and assembled into plasmids via Gibson Assembly using the NEBuilder HiFi DNA Assembly kit (New England Biosciences). Vectors pSBS4 (empty vector), pJLG091 (P_slp_–*Pf*αgal), and pJLG093 (P_slp_–*Cm*βgal) were then cloned into chemically competent *E. coli* 10beta, isolated using ZymoPURE miniprep kits (Zymo Research), and sequence confirmed (Azenta Genewiz). pJLG161 is identical to pJLG093 except that expression of *Cm*βgal is driven instead by P_bh_ (**Fig. 1**). pJLG161 was constructed from pJLG093 in partnership with the Department of Energy Joint Genome Institute (JGI) at Lawrence Berkely National Lab (Berkely, CA) as described below. pJLG093 was first modified to create unique PmeI sites, aiding subsequent promoter insertion. The vector was linearized by PCR amplification (**Table S1;** Primers B431.093.VM.F & VM.R), and re-circularized via Gibson assembly together with an ultramer (**Table S1;** Ultramer JGI.UM1) purchased from Integrated DNA Technologies, using the NEBuilder HiFi DNA assembly kit. After validation of the modified vector, the sequence corresponding to P_bh_ was flanked by linkers designed for assembly into pJLG093_PmeI linearized by PmeI digest (**Table S2**), purchased (Twist Biosciences) and assembled using the NEBuilder HiFi kit. These assemblies were subsequently transformed into chemically competent *E. coli* Top10 of which candidate colonies were picked, sequence verified on the Pacific Biosciences Revio platform (Pacific Biosciences), and analyzed using custom pipelines at the Joint Genome Institute. pJLG161 was subsequently isolated using ZymoPURE miniprep kits (Zymo Research), and sequence confirmed (Azenta Genewiz).

**Figure 1.**
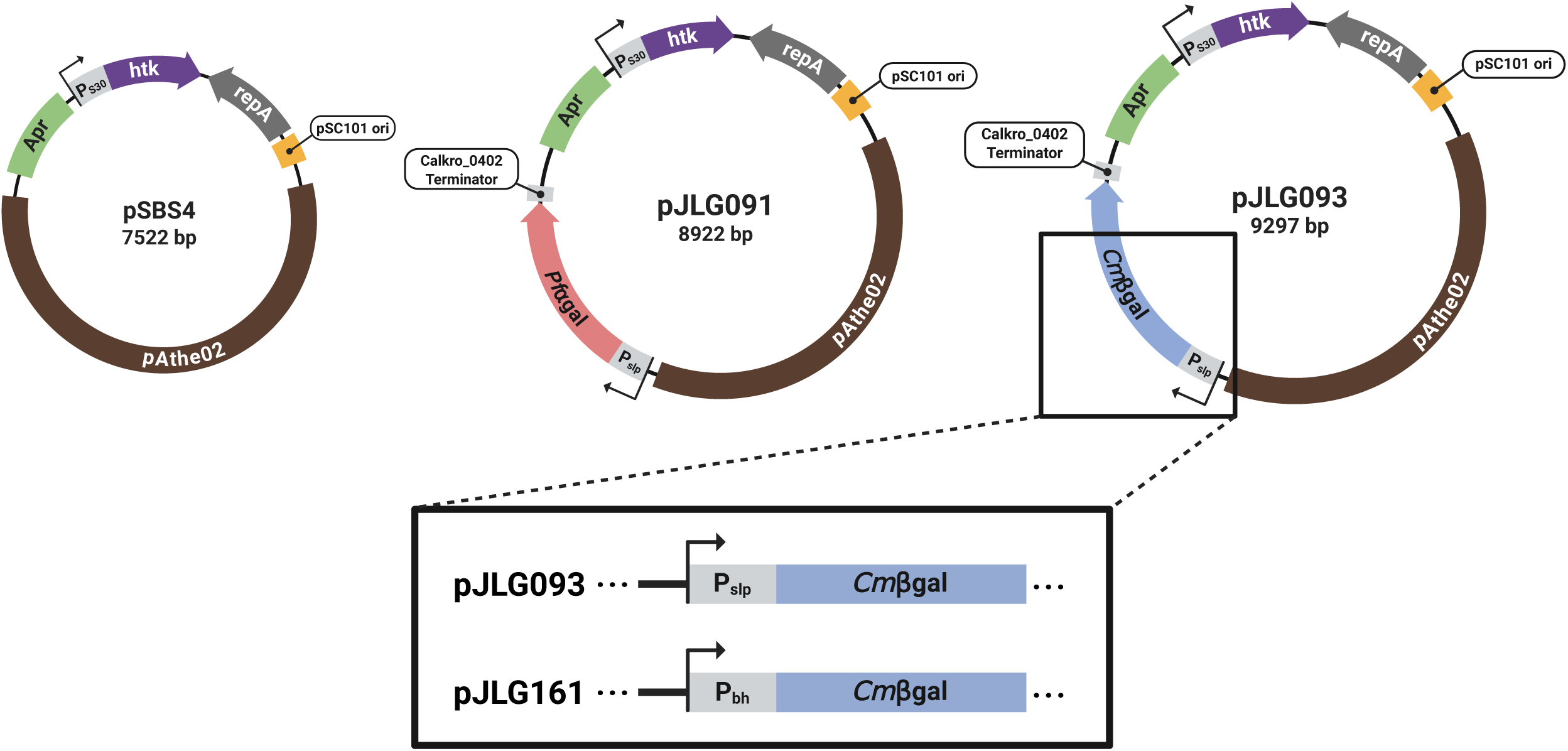
Maps of plasmids constructed and transformed into *A. bescii*. pSBS4 (Empty Vector) was used for the original implementation of the highly thermostable kanamycin (*htk*) selection marker in *A. bescii* (Lipscomb *et al*., 2016). This vector contains a native *A. bescii* replicating plasmid (pAthe02) which provides machinery for replication in *A. bescii*, the *htk* gene expressed by P_S30_ for selection on kanamycin, as well as elements for cloning in *E. coli* including: an apramycin resistance marker (Apr), replication initiation protein A (repA), and the pSC101 origin. pSBS4 was modified to add a reporter expression site between Apr and pAthe02, resulting in: pJLG091 (P_slp_-*Pf*αgal), pJLG093 (P_slp_-*Cm*βgal), and pJLG161 (P_bh_-*Cm*βgal). Created in BioRender. Galindo, J. (2025) https://BioRender.com/drjomi3.

**Table 1.**
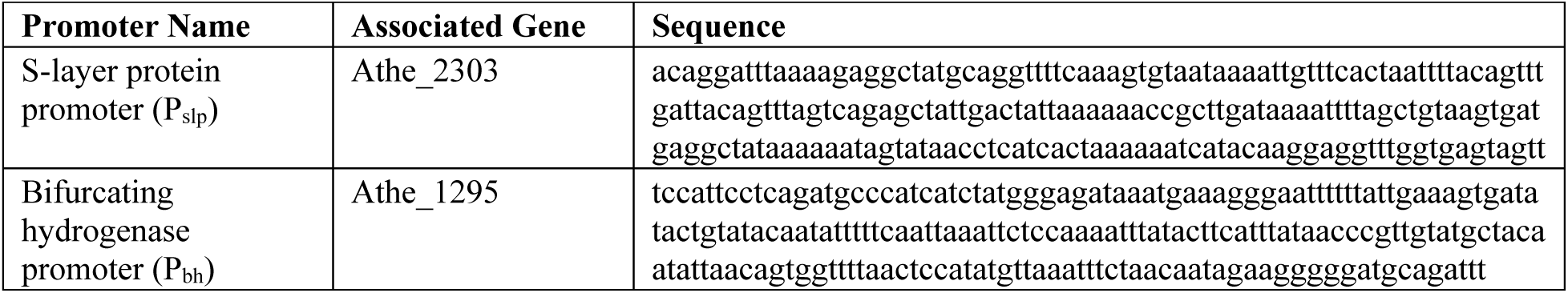
Promoter sequences used to drive galactosidase reporter expression in *A. bescii*.

### Plasmid preparation and transformation into *A. bescii*

For transformation in *A. bescii*, larger quantities of plasmid DNA were extracted from *E. coli* using the ZymoPURE maxiprep kit (Zymo Research). Extracted plasmids were then methylated *in vitro* using the M.CbeI methyltransferase and purified via phenol-chloroform extraction as previously described (Chung *et al*., 2012; Lipscomb *et al*., 2016). Wild type *A. bescii* DSM 6725 was obtained from the lab of Dr. Robert Kelly (North Carolina State University). Competent *A. bescii* were grown on CM516 media containing amino acids (CM516-AA) to an optical density at 680 nm (OD680) of 0.04-0.08 and prepared for transformation as described previously (Lipscomb *et al*., 2016). 50 µl of competent cells were transformed with 1-2 µg of plasmid in a 1mm gap electroporation cuvette using a Bio-Rad gene pulser at 1800 V, 400Ω, and 25 µF. Electroporated cells were immediately resuspended in 1 ml of CM516 media and transferred to 10 ml tubes containing the same media pre-heated to 70 °C. Cells were allowed to recover for 90 minutes before being transferred to pre-heated bottles containing 50 ml of selective CM516K media. After 24-36 hours of growth, cells were passaged into 10 ml of fresh CM516K media and allowed to grow overnight. Passaged cells were then plated and grown for 48 hours in solid selective CM516K media with 1.5% (w/v) agar at 70 °C under a 95% (v/v) N_2_ and 5% (v/v) H_2_ atmosphere in an anaerobic chamber. Single colonies were picked and screened via colony PCR (**Fig. S1**) using primers JLG181 and JLG224 (**Table S1**). Presence of the correct promoter-reporter sequences were confirmed by long-read sequencing of colony PCR products (Azenta Genewiz, PCR-EZ).

### Enzymatic reporter assay

To detect galactosidase activity in *A. bescii* cells, para-nitrophenol-α-D-galactopyranoside (pNPαGal) and para-nitrophenol-β-D-galactopyranoside (pNPβGal) obtained from TCI chemicals, were used as colorimetric substrates. Substrate solutions contained 5 mM pNPαGal or pNPβGal dissolved in 100 mM sodium acetate pH 5.5 buffer which was chosen based on the previously determined optimal pH ranges of *Pf*αgal and *Cm*βgal (van Lieshout *et al*., 2003; Letsididi *et al*., 2017). *A. bescii* cells were prepared for galactosidase assays by first pelleting 5-15 ml of freshly grown cells at the maximum rotor speed (7000 x g for 15 ml pellets or 21000 x g for 5 ml pellets) for 10 minutes, followed by removal of the supernatant and storage at −80 °C for later use. Immediately prior to testing, cell pellets were resuspended and concentrated 3-5x in 1-3 ml of 100 mM pH 5.5 sodium acetate buffer to a final OD680 of 0.35-0.5, measured on a Nanodrop One C spectrophotometer with 100 mM sodium acetate buffer as the blanking solution. For assays involving heat-treatments, 50-100 µl of cells or blank buffer were aliquoted into PCR strip tubes and incubated in a thermocycler at 90 or 98 °C for 10 minutes unless described otherwise. To begin the reaction, 10-30 µL of cells or blank buffer were added to 60-80 µL of substrate solution or blank buffer to a total volume of 90 µL. Assays that involved wild type or the P_slp_–*Pf*αgal strain of *A. bescii* required 30 µl of cells, while testing of *Cm*βgal expressing *A. bescii* only required 10 µl of cells per reaction. Reactions were incubated in a thermocycler at the appropriate temperature for the experimentally prescribed time after which all reactions were immediately quenched with the addition of 180 µL of 1M sodium carbonate. The absorbance at 405 nm (A405) of 100 µl of each reaction was then measured in a flat-bottomed clear 96 well plate using a BioTek SynergyH1 microplate reader (Agilent). For all reaction conditions the following controls were included: a substrate only (no cell) condition to account for the thermal background degradation of substrate, a no substrate condition for each cell type to account for background scattering from cellular debris, and a buffer only condition to isolate the absorbance due to debris in the prior control from the buffer itself. All reaction conditions were performed in technical triplicate.

Normalized galactosidase activity was calculated as defined in equation (1) based on the equations in “Experiments in Molecular Genetics” for measuring β-galactosidase activity in *E. coli* using o-nitrophenyl-beta-D-galactopyranoside (Miller, 1972). The most notable modifications to are cellular debris background is explicitly accounted for with a series of control reactions rather than estimated with the absorbance at 550 nm, and normalization is done with the optical density at 680 nm (OD680) rather than that at 600 nm (OD600). In equation (1), the A405 of the no cell control (A405_NC_) is subtracted from the A405 of the experimental condition (A405_exp_) to remove thermal background degradation of substrate. Separately, the A405 of the buffer only control (A405_BO_) is subtracted from that of the no substrate control (A405_NS_). This is then subtracted from the A405_exp_ - A405_NS_ difference to account for debris scattering. This final value is then divided by the previously measured OD680 of the resuspended *A. bescii* input to the assay to normalize for differences in the amount of cells added.

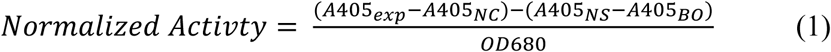

### Assessment of *Pfα*gal and *Cm*βgal as reporters in *A. bescii*

To test for background activity from endogenous galactosidases, wild type *A. bescii* DSM 6725 were grown on CM516 media to an OD680 of 0.15 (late exponential), pelleted, and frozen. Cells were resuspended and heat-treated at 90 or 98 °C for 0, 10, 30, or 60 minutes prior to adding pNPαGal or pNPβGal solutions in the enzyme assay described above, with incubation for 2 hours at 75 °C.

Prior to initial testing of the hyperthermophilic galactosidase reporters, *A. bescii* containing the Empty Vector, P_slp_–*Pf*αgal, and P_slp_–*Cm*βgal, were grown on selective CM516K media to an OD680 of 0.1-0.12 (mid-late exponential) and harvested as described previously. For time course experiments, cells that contained the Empty Vector or P_slp_–*Pf*αgal were prepared and heat-treated at 98 °C. Cells were then added to pNPαGal solution and incubated for 0, 1, 2, or 3 hours at 98 °C. Separately this was repeated for Empty Vector and P_slp_–*Cm*βgal cells except heat-treatments were carried out at 90 °C, cells were instead added to pNPβGal solution, and incubations were carried out at 90 °C for 0, 10, 20, or 30 minutes. To test the effects of various heat-treatment conditions on the reporters, resuspended Empty Vector, P_slp_–*Pf*αgal, and P_slp_–*Cm*βgal cells were heat-treated at 90 or 98 °C which were subsequently added alongside un-heat-treated cells to both the pNP substrate solutions. For assays testing P_slp_–*Pf*αgal against the Empty Vector, cells were incubated for 2 hours, while for assays testing P_slp_–*Cm*βgal, cells were only incubated for 20 minutes. Signal to noise ratio was calculated as the normalized activity of reporter expressing cells on their respective preferred pNP substrates divided by that of the empty vector control at the corresponding conditions (i.e. P_slp_–*Pf*αgal/Empty Vector activity on pNPαGal, or P_slp_–*Cm*βgal/Empty Vector activity on pNPβGal).

### Testing *Cm*βgal activity throughout the growth of *A. bescii*

To start the growth curves of *A. bescii*, strains containing pSBS4 (empty vector), pJLG093 (P_slp_–*Cm*βgal), and pJLG161 (P_bh_–*Cm*βgal) were inoculated at a target OD680 of 0.002 in 50 mL of selective CM516K media. Cultures were grown for 29 hours in biological triplicate at 70 °C, with each culture’s OD680 measured at intervals of roughly 3-5 hours. At 12, 18, 24, and 29 hours, 4-5 ml of each culture was removed, after which cells were pelleted, frozen, and assayed as described above. To test for the activity of *Cm*βgal, thawed pellets were prepared as described above with heat treatment at 90 °C. Cells were then added to pNPβGal solution and incubated for 10 minutes at 90 °C.

### RNA extraction and qRT-PCR of *Cm*βgal expressing *A. bescii*

Similar to previous studies, RNA was isolated from *A. bescii* containing pSBS4, pJLG093, and pJLG161 that were grown on CM516K media for 18 hours to OD680 values of 0.11-0.16 (mid-late exponential phase), with three biological replicates for each strain (Williams-Rhaesa *et al*., 2018; Straub *et al*., 2020; Tanwee *et al*., 2023; Bing *et al*., 2024). After growth, 30-40 ml of cells were immediately pelleted at 6000 x g for 10 minutes and, after removal of the supernatant, frozen at −80 °C. Prior to purification, thawed cell pellets were lysed as previously described with the addition of 240 µl of cold PBS, 75 µl of lysozyme (20 mg/ml), and 300 µl of the Monarch® gDNA Tissue Lysis Buffer (New England Biosciences), followed by incubation at 37°C for 15 minutes (Bing *et al*., 2024). 300 µl lysate from each pellet was then added to two volumes (600 µl) of Monarch® StabiLyse DNA/RNA Buffer (New England Biosciences) (Bing *et al*., 2024). From this, RNA was purified using the Monarch^®^ Spin RNA Isolation Kit (New England Biosciences) as per the manufacturer’s instructions with the on-column DNase I treatment step. RNA concentrations were quantified using a Nanodrop One spectrophotometer (Thermo Scientific). qRT-PCR assays were carried out on a Viia7^TM^ Real-Time PCR System (Thermo Scientific). qRT-PCR on extracted RNA samples was performed using the Luna^®^ Universal One-Step RT-qPCR Kit (New England Biosciences) according to the manufacturer protocol, with 50 ng of total RNA added to 10 µl reactions in a 384 well plate. A no-RT control condition was included for each experimental condition to check for DNA contamination. All reaction conditions, including for each biological replicate, were performed in technical triplicate. Expression of the *Cmβgal* gene (**Table S1;** Primers JLG219-220) was calculated relative to that of the *A. bescii gapdh* (*Athe_1406*) using primers (**Table S1;** Primers JLG211_CTS480-JLG212_CTS481) utilized in a previous *A. bescii* study (Straub *et al*., 2020).

## Results

### Implementation of two hyperthermophilic galactosidases as reporters in *A. bescii*

The genome of wild type *A. bescii* contains least one characterized α-galactosidase, along with several putative α- and β-galactosidases (Lee *et al*., 2017; Drula *et al*., 2022). To assess the level of heat-treatment needed to eliminate background activity from these enzymes on colorimetric pNP-glycoside substrates, prepared wild type *A. bescii* cells were heat-treated at 90 or 98 °C for 0-60 minutes. Cells were then added to solutions of pNPαGal or pNPβGal and incubated for 2 hours at 75 °C to test for endogenous α- or β-galactosidase activity respectively. Significant background activity was detected on both of these substrates with *A. bescii* that were not heat-treated (**Fig. 2**). However, this background activity was eliminated by heat-treatment for as short as 10 minutes at either 90 or 98 °C, indicating native *A. bescii* α- and β-galactosidase enzymes are inactivated with this relatively short incubation at temperatures above the native growth temperature of *A. bescii* (**Fig. 2**).

**Figure 2.**
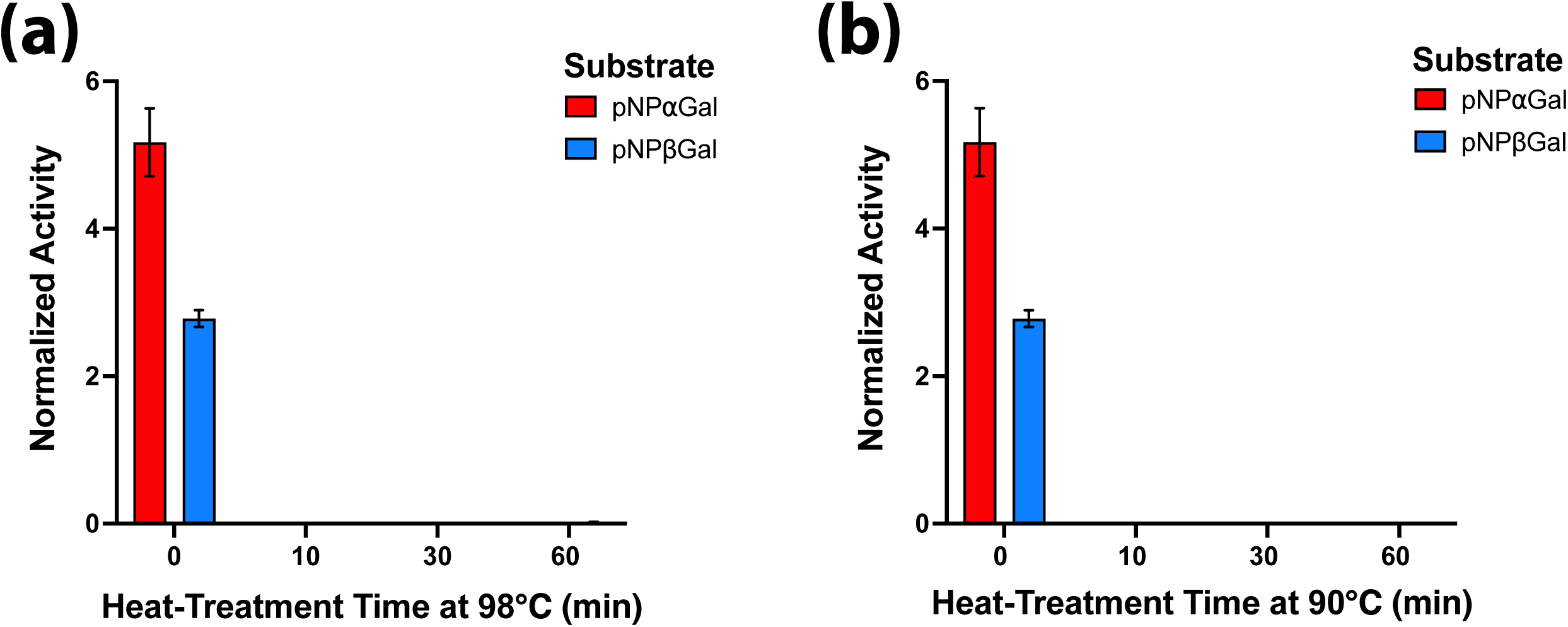
Native galactosidase activity of *A. bescii* is eliminated by heat-treatment. Activity from wild type *A. bescii* DSM 6725 cells on pNPαGal (red) or pNPβGal (blue) as measured in a two-hour assay at 75 °C after being heat treated at: **(a)** 98 °C for 0, 10, 30 or 60 minutes; **(b)** 90 °C for 0, 10, 30 or 60 minutes. Error bars represent one standard deviation between triplicate technical replicates at each reaction condition.

To determine if *Pf*αgal and *Cm*βgal could serve as effective reporters, wild type *A. bescii* DSM 6725 was transformed with plasmids pJLG091 (P_slp_–*Pf*αgal) and pJLG093 (P_slp_–*Cm*βgal), which drive strong constitutive expression of the galactosidases with P_slp_, as well as the plasmid pSBS4 (Empty Vector) to serve as an empty vector control (**Fig. 1; Table 1**). Previous characterization efforts describe both enzymes as having optimal activity at temperatures >100 °C, however these studies also show that only *Pf*αgal is stable for long periods at these temperatures while *Cm*βgal begins to lose activity rapidly within 90 minutes at temperatures >90 °C (van Lieshout *et al*., 2003; Letsididi *et al*., 2017). Thus, the ability of both reporters to function at 90 or 98 °C, and the necessity of heat-treatment given the elevated reaction temperatures, were evaluated.

Empty Vector, P_slp_–*Pf*αgal, and P_slp_–*Cm*βgal cells were heat-treated at 90 or 98 °C and tested alongside un-heat-treated cells at the same temperatures on both pNPαGal and pNPβGal. These results showed that both systems produced clear signals above the background observed in the empty vector control (**Fig. 3a-h**). However, to produce a sufficiently strong signal while maintaining approximately linear behavior with respect to incubation time, P_slp_–*Pf*αgal cells required incubations with substrate for at least 2-3 hours, while ideal reaction times for P_slp_– *Cm*βgal cells were only 10-20 minutes (**Fig. 3a & b**). These results also show that heat-treatments significantly reduce background for both reporters (**Fig. 3c-h**). *Cm*βgal in particular shows a marked improvement in signal to noise ratio, increasing from 32.2x to 170.4x background with heat-treatment (**Fig. 3h**). As expected, *Pf*αgal is functional at both 90 and 98 °C (**Fig. 3c & d)**. In general, *Pf*αgal produces a ∼10x weaker signal compared with that of *Cm*βgal, with the highest signal to noise ratio observed in this test of 12.4x background (**Fig. 3e**). *Cm*βgal on the other hand appears to be briefly active at 98 °C when no heat-treatment step is included, but 10 minutes of heat-treatment at 98 °C appears to eliminate activity from this enzyme (**Fig. 3f & g**). Finally, testing of these galactosidase expressing cells on both pNPαGal and pNPβGal substrates show these reporters act orthogonally to each other, with no activity above that of the empty vector control detected on their non-preferred substrate (**Fig. S2a-d**). After this testing, *Cm*βgal was chosen for subsequent tests of expression in *A. bescii* as it produced a far higher signal to noise ratio while requiring much shorter incubations compared with *Pf*αgal.

**Figure 3.**
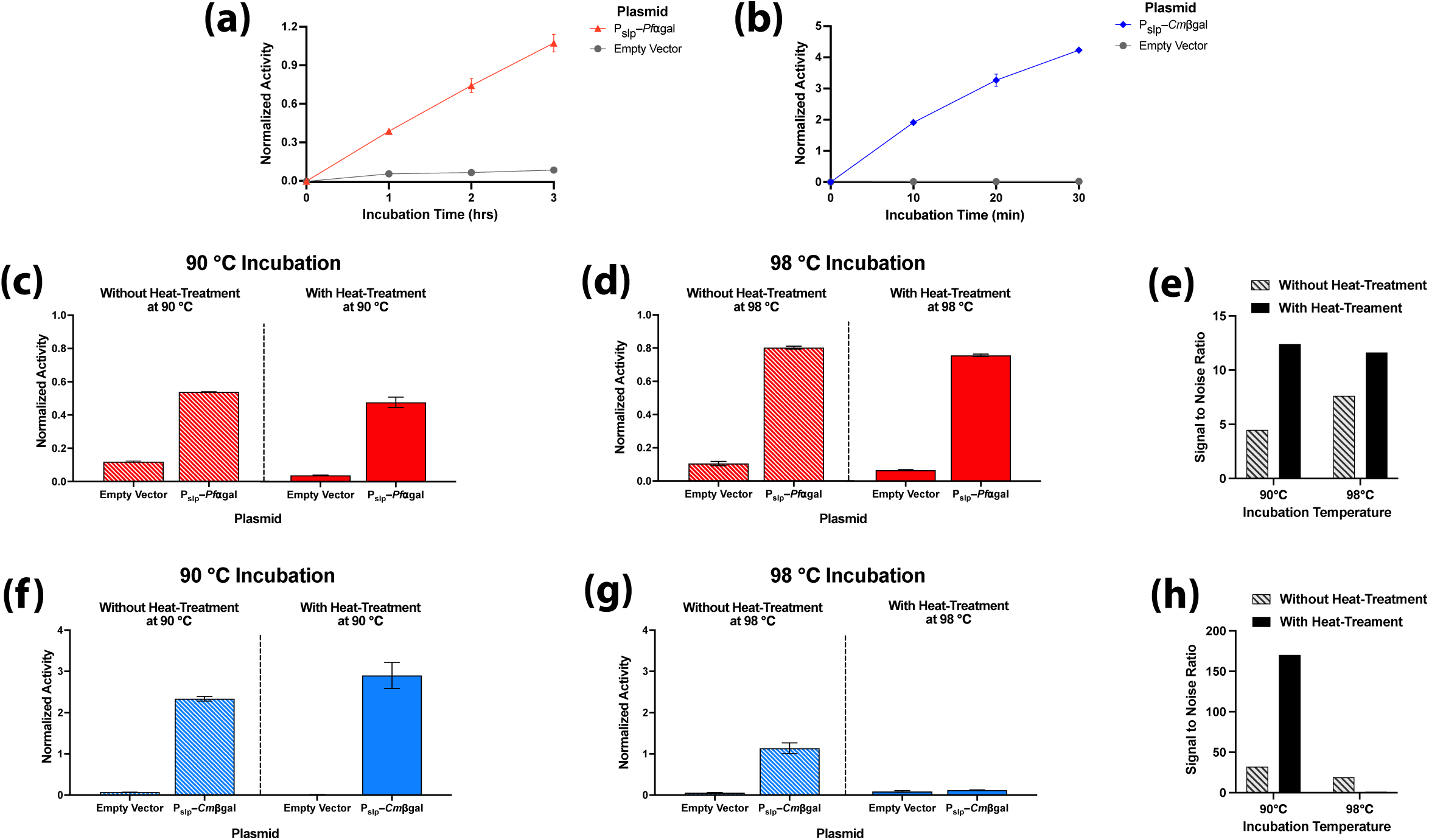
Development of reaction conditions for assessing expression of the *Pf*αgal and *Cm*βgal as hyperthermophilic galactosidase reporters in *A .bescii.* Here expression of both reporters is driven by P_slp_. Activity detected after various incubation times with pNP substrate compared with the Empty Vector strain for: **(a)** *Pf*αgal on pNPαGal for 0-3 hours at 98 °C; **(b)** *Cm*βgal on pNPβGal for 0-30 minutes 90 °C. Activity of *Pf*αgal vs. the Empty Vector strain on pNPαGal for: **(c)** 2 hours at 90 °C with and without 10 minutes of heat-treatment at 90 °C; **(d)** 2 hours at 98 °C with and without 10 minutes of heat-treatment at 98 °C; **(e)** The signal to noise ratio of *Pf*αgal on pNPαGal defined as the activity of *Pf*αgal divided by that of the Empty Vector strain after each of the four incubation conditions in **(c)** and **(d)**. Activity of *Cm*βgal vs. the Empty Vector strain on pNPβGal for: **(f)** 20 minutes at 90 °C with and without 10 minutes of heat-treatment at 90 °C; **(g)** 20 minutes at 98 °C with and without 10 minutes of heat-treatment at 98°C; **(h)** The signal to noise ratio of *Cm*βgal on pNPβGal defined as the activity of *Cm*βgal divided by that of the Empty Vector strain after each of the four incubation conditions in **(f)** and **(g)**. Error bars in all panels represent one standard deviation calculated from triplicate technical replicates at each reaction condition.

### Utilizing *Cm*βgal to distinguish differences in protein expression in *A. bescii*

Next, reporter expression was tested over the growth of *A. bescii* strains containing the *Cm*βgal reporter under the control of two previously utilized promoters, P_slp_ and P_bh_. Based on previous studies, the P_bh_ promoter should drive somewhat lower expression than P_slp_ (Williams-Rhaesa *et al*., 2018). *A. bescii* strains containing pSBS4 (Empty Vector), pJLG093 (P_slp_–*Cm*βgal), and pJLG161 (P_bh_–*Cm*βgal) were grown and monitored over the course of twenty-nine hours in biological triplicate (**Fig. 4a**). At time points of 12, 18, 24, and 29 hours, roughly corresponding to exponential, late exponential, early stationary, and stationary phase growth, respectively, cells were harvested for enzyme reporter measurement on pNPβGal with heat-treatment and incubation at 90 °C (**Figure 4b**). As expected, no significant activity was detected from the Empty Vector strain at any stage of growth (**Fig. 4b**). These results also show that the relative strength of enzyme activity from P_bh_–*Cm*βgal to that of P_slp_–*Cm*βgal changes significantly throughout growth, with relative activities of 37%, 56%, 72%, and 73% at 12, 18, 24, and 29 hours of growth, respectively (**Fig. 4b**). In general, activity from both promoters appears to increase as *A. bescii* enters stationary phase (24- and 29-hour timepoints), though variability in expression between biological replicates also increases in stationary phase (**Fig. 4b**).

**Figure 4.**
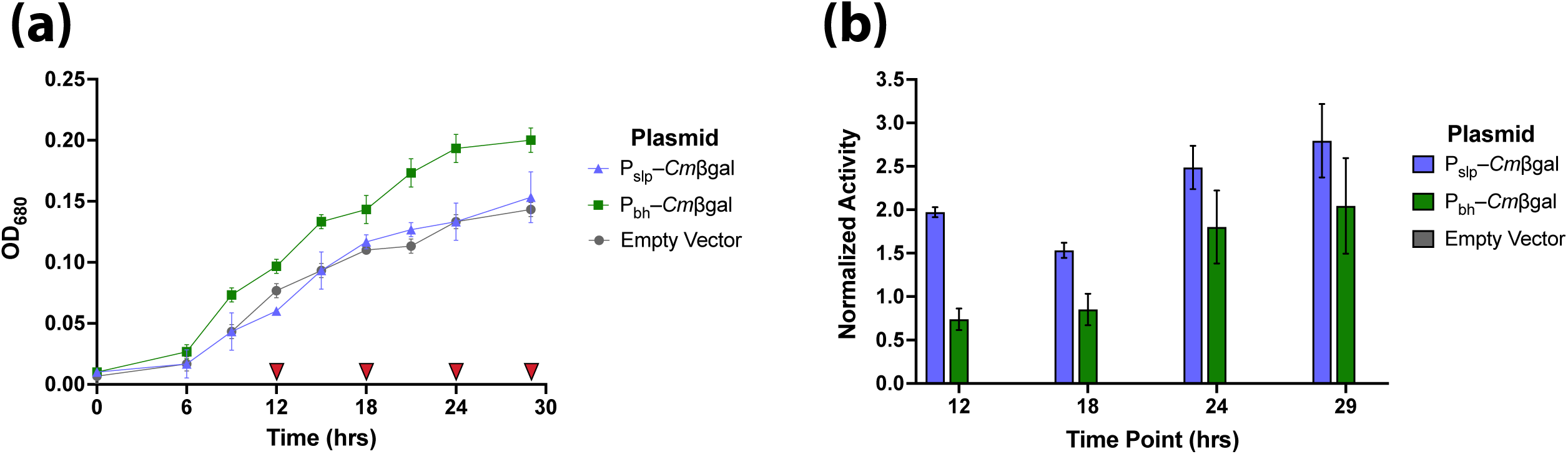
**(a)** Growth of *A. bescii* strains transformed with plasmids expressing *Cm*βgal with P_slp_ or P_bh_ as well as the Empty Vector strain over the course of 29 hours. Red triangles (▾) indicate time points (12, 18, 24, and 29 hours) where cells were harvested for enzyme assays. **(b)** Corresponding normalized β-galactosidase activity of prepared *A. bescii* cells at 12, 18, 24, and 29 hours. Cells were heat-treated for 10 minutes at 90 °C followed by another 10-minute incubation at 90 °C on pNPβgal. Error bars in both **(a)** and **(b)** represent one standard deviation between triplicate biological replicates.

To assess how *Cm*βgal reporter activity levels mirror transcription levels, qRT-PCR was performed on the *Cmβgal* gene. RNA was extracted from empty vector, P_slp_–*Cm*βgal, and P_bh_– *Cm*βgal *A. bescii* strains in late exponential phase (18-hour timepoint) grown in biological triplicate. Levels of *Cm*βgal transcription in each strain were calculated relative to that of the endogenous *A. bescii* glyceraldehyde-3-phosphate dehydrogenase *gapdh* (*Athe_1406*) housekeeping gene as is standard in the literature (Williams-Rhaesa *et al*., 2018; Straub *et al*., 2020; Tanwee *et al*., 2023). Results show that both P_slp_ and P_bh_ drive strong levels of transcription, with expression of 15.8x and 4.8x that of *gapdh,* respectively (**Fig. 5**). P_slp_ is the stronger promoter with an average level of transcription 3.3x that of P_bh_ (**Fig. 5**). This mirrors a smaller difference in enzyme activity, where *Cm*βgal expressed by P_slp_ produced an average activity 1.8x that driven by P_bh_ (**Fig. 4b**).

**Figure 5.**
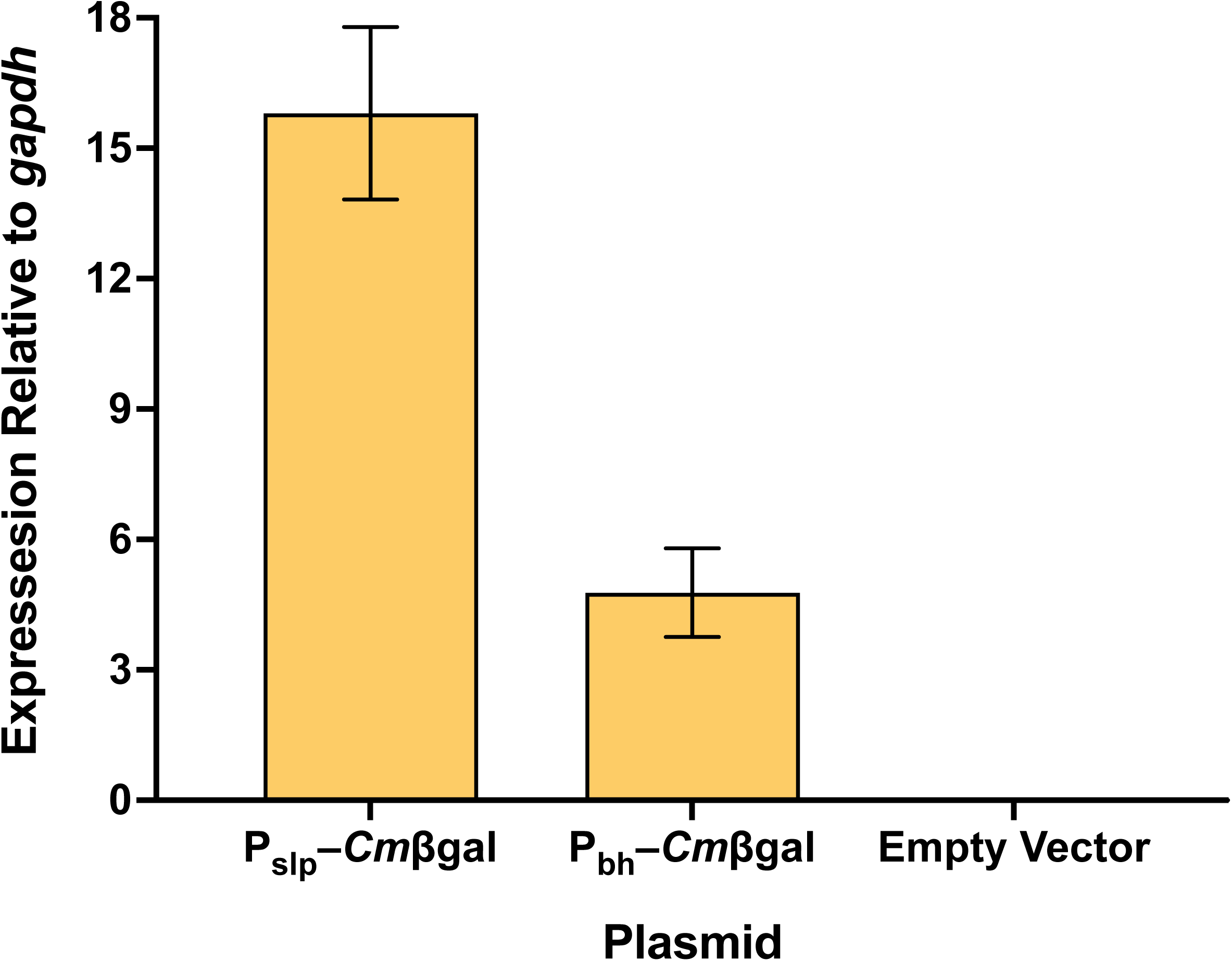
Transcription of the *Cmβgal* gene in transformed *A. bescii* grown for 18 hours relative to that of the endogenous *gapdh* (*Athe_1406*) housekeeping gene. Error bars represent one standard deviation between biological triplicates.

## Discussion

In this work we demonstrate two hyperthermophilic galactosidases, *Pf*αgal and *Cm*βgal, can be used as enzymatic reporters in *A. bescii* to robustly report enzyme expression driven by different promoters (**Fig. 3**). Heat-treatment for as short as 10 minutes at 90 °C is sufficient to eliminate any activity from endogenous *A. bescii* galactosidases (**Fig. 2**), reducing background and improving reporter signal to noise (**Fig. 3**). These reporters function orthogonally, with no activity on their non-preferred pNP substrate (**Fig. S2**) opening the possibility that dual expression within a single strain could be used to produce distinct readouts. This could prove useful for certain applications such as engineering transcriptional terminators. Of these two reporter enzymes, *Cm*βgal is the stronger reporter with a far greater dynamic range in shorter assay time. While *Pf*αgal requires incubations on the order of several hours at 98 °C, *Cm*βgal produces ∼10x the signal of *Pf*αgal with less cells in as short as 10 minutes of incubation at 90 °C (**Fig. 3a & b**).

By expressing *Cm*βgal with two different promoters, P_slp_ and P_bh_, we show that our hyperthermophilic β-galactosidase based system can robustly measure differential enzymatic expression in *A. bescii* (**Fig. 4a & b**). The strength of both promoters appears to increase but become more variable as cells enter stationary phase (**Fig. 4b**). While P_slp_ consistently drives higher expression, P_bh_ does seem to increase in relative strength in stationary phase (**Fig. 4b**). Finally, through qRT-PCR we show that at the transcriptional level P_slp_ produces RNA expression greater than three times that of P_bh_ (**Fig. 5**). This is consistent with the previous observations of Williams-Rhaesa *et al*. who report that for chromosomally integrated constructs P_slp_ drives transcription at levels between 3-6x that of P_bh_ (Williams-Rhaesa *et al*., 2018). This suggests that *Cm*βgal activity as measured by the reporter assay can serve as a representative measure of transcription, however relative enzymatic activity may not perfectly align to that of mRNA concentration. The results in this case would suggest that a higher level of transcription may only produce a marginal increase in observed galactosidase activity. This discrepancy could be accounted by a difference in the strengths of the ribosome binding sites in P_slp_ and P_bh_ which could significantly alter the rate protein translation between the two promoters (Kent and Dixon, 2020).

Taken as a whole, we describe an easy to implement and robust enzymatic reporter system in *A. bescii* that could serve a variety of purposes from genetic tool development to strain identification. While demonstrated here in *A. bescii*, we expect this hyperthermophilic enzyme reporter system could easily be adapted for use in other thermophilic anaerobic species and would be especially valuable in those which grow at temperatures >70°C where other anaerobic reporters are not viable, or those where native glycosidase enzymes obscure less thermophilic enzymatic reporters. Ultimately this reporter system will enable the development of new genetic tools, metabolic engineering approaches, and next generation bioprocessing efforts using anaerobic thermophiles.

## Author Contribution

**Joey L. Galindo:** Conceptualization, Methodology, Formal analysis, Investigation, Visualization, Writing - Original Draft

**Hansen Tjo:** Formal analysis, Investigation, Writing - Review & Editing

**Jonathan M. Conway:** Conceptualization, Supervision, Resources, Writing - Review & Editing, Funding acquisition

## Acknowledgements

The construction of pJLG161 was conducted by the Joint Genome Institute (https://ror.org/04xm1d337) under proposal: **10.46936/10.25585/60012765**. The U.S. Department of Energy Joint Genome Institute, a DOE Office of Science User Facility, is supported by the Office of Science of the U.S. Department of Energy operated under Contract No. DE-AC02-05CH11231. H.T. acknowledges the support of the William Clay Ford, Jr ‘79 and Lisa Vanderzee Ford ‘82 Graduate Fellowship Fund administered by the High Meadows Environmental Institute at Princeton University.

## References

Bing, R.G. et al. (2021) ‘Thermophilic microbial deconstruction and conversion of natural and transgenic lignocellulose’, Environmental Microbiology Reports, 13(3), pp. 272–293. Available at: 10.1111/1758-2229.12943.

Bing, R.G. et al. (2024) ‘Engineering ethanologenicity into the extremely thermophilic bacterium *Anaerocellum (*f. *Caldicellulosiriuptor) bescii*’, Metabolic Engineering, 86, pp. 99–114. Available at: 10.1016/j.ymben.2024.09.007.

Blumer-Schuette, S.E. et al. (2014) ‘Thermophilic lignocellulose deconstruction’, FEMS Microbiology Reviews, 38(3), pp. 393–448. Available at: 10.1111/1574-6976.12044.

Blumer-Schuette, S.E. (2020) ‘Insights into Thermophilic Plant Biomass Hydrolysis from *Caldicellulosiruptor* Systems Biology’, Microorganisms, 8(3), p. 385. Available at: 10.3390/microorganisms8030385.

Cha, M. et al. (2013) ‘Metabolic engineering of *Caldicellulosiruptor bescii* yields increased hydrogen production from lignocellulosic biomass’, Biotechnology for Biofuels, 6(1), p. 85. Available at: 10.1186/1754-6834-6-85.

Chung, D. et al. (2012) ‘Methylation by a Unique α-class N4-Cytosine Methyltransferase Is Required for DNA Transformation of *Caldicellulosiruptor bescii* DSM6725’, PLOS ONE, 7(8), p. e43844. Available at: 10.1371/journal.pone.0043844.

Chung, D. et al. (2013) ‘Construction of a Stable Replicating Shuttle Vector for *Caldicellulosiruptor* Species: Use for Extending Genetic Methodologies to Other Members of This Genus’, PLOS ONE, 8(5), p. e62881. Available at: 10.1371/journal.pone.0062881.

Conway, J.M. et al. (2018) ‘Parsing in vivo and in vitro contributions to microcrystalline cellulose hydrolysis by multidomain glycoside hydrolases in the *Caldicellulosiruptor bescii* secretome’, Biotechnology and Bioengineering, 115(10), pp. 2426–2440. Available at: 10.1002/bit.26773.

Drula, E. et al. (2022) ‘The carbohydrate-active enzyme database: functions and literature’, Nucleic Acids Research, 50(D1), pp. D571–D577. Available at: 10.1093/nar/gkab1045.

Fujita, A. et al. (2015) ‘A reporter gene system for the precise measurement of promoter activity in *Thermus thermophilus* HB27’, Extremophiles, 19(6), pp. 1193–1201. Available at: 10.1007/s00792-015-0789-3.

Ganguly, J., Martin-Pascual, M. and van Kranenburg, R. (2020) ‘CRISPR interference (CRISPRi) as transcriptional repression tool for *Hungateiclostridium thermocellum* DSM 1313’, Microbial Biotechnology, 13(2), pp. 339–349. Available at: 10.1111/1751-7915.13516.

Hocq, R. et al. (2023) ‘A fluorescent reporter system for anaerobic thermophiles’, Frontiers in Bioengineering and Biotechnology, 11. Available at: 10.3389/fbioe.2023.1226889.

Honarbakhsh, M. et al. (2012) ‘Development of a thermostable β-glucuronidase-based reporter system for monitoring gene expression in hyperthermophiles’, Biotechnology and Bioengineering, 109(7), pp. 1881–1886. Available at: 10.1002/bit.24432.

Jensen, T.Ø. et al. (2017) ‘Application of the thermostable β-galactosidase, BgaB, from *Geobacillus stearothermophilus* as a versatile reporter under anaerobic and aerobic conditions’, AMB Express, 7(1), p. 169. Available at: 10.1186/s13568-017-0469-z.

Kent, R. and Dixon, N. (2020) ‘Contemporary Tools for Regulating Gene Expression in Bacteria’, Trends in Biotechnology, 38(3), pp. 316–333. Available at: 10.1016/j.tibtech.2019.09.007.

Kim, N.M., Sinnott, R.W. and Sandoval, N.R. (2020) ‘Transcription factor-based biosensors and inducible systems in non-model bacteria: current progress and future directions’, Current Opinion in Biotechnology, 64, pp. 39–46. Available at: 10.1016/j.copbio.2019.09.009.

Langholtz, M. et al. (2024) 2023 *Billion-Ton Report: An Assessment of U.S. Renewable Carbon Resources*. ORNL/SPR-2024/3103, 2441098, p. ORNL/SPR-2024/3103, 2441098. Available at: 10.2172/2441098.

Lee, A. et al. (2017) ‘Characterization of a thermostable glycoside hydrolase family 36 α-galactosidase from *Caldicellulosiruptor bescii*’, Journal of Bioscience and Bioengineering, 124(3), pp. 289–295. Available at: 10.1016/j.jbiosc.2017.04.011.

Lee, L.L. et al. (2020) ‘The biology and biotechnology of the genus *Caldicellulosiruptor*: recent developments in “Caldi World”’, Extremophiles, 24(1), pp. 1–15. Available at: 10.1007/s00792-019-01116-5.

Letsididi, R. et al. (2017) ‘Characterization of a thermostable glycoside hydrolase (CMbg0408) from the hyperthermophilic archaeon *Caldivirga maquilingensis* IC-167’, Journal of the Science of Food and Agriculture, 97(7), pp. 2132–2140. Available at: 10.1002/jsfa.8019.

van Lieshout, J.F.T. et al. (2003) ‘Identification and Molecular Characterization of a Novel Type of α-galactosidase from *Pyrococcus furiosus*’, Biocatalysis and Biotransformation, 21(4–5), pp. 243–252. Available at: 10.1080/10242420310001614342.

Lipscomb, G.L., et al. (2016) ‘A Highly Thermostable Kanamycin Resistance Marker Expands the Tool Kit for Genetic Manipulation of *Caldicellulosiruptor bescii*’, Applied and Environmental Microbiology. Edited by V. Müller, 82(14), pp. 4421–4428. Available at: 10.1128/AEM.00570-16.

Loder, A.J. et al. (2017) ‘Extreme Thermophiles as Metabolic Engineering Platforms: Strategies and Current Perspective’, in *Industrial Biotechnology*. John Wiley & Sons, Ltd, pp. 505–580. Available at: 10.1002/9783527807796.ch14.

Lynd, L.R. et al. (2022) ‘Toward low-cost biological and hybrid biological/catalytic conversion of cellulosic biomass to fuels’, Energy & Environmental Science, 15(3), pp. 938–990. Available at: 10.1039/D1EE02540F.

Mattossovich, R. et al. (2020) ‘A journey down to hell: new thermostable protein-tags for biotechnology at high temperatures’, Extremophiles, 24(1), pp. 81–91. Available at: 10.1007/s00792-019-01134-3.

Merlo, R. et al. (2022) ‘First thermostable CLIP-*tag* by rational design applied to an archaeal *O6*-alkyl-guanine-DNA-alkyl-transferase’, Computational and Structural Biotechnology Journal, 20, pp. 5275–5286. Available at: 10.1016/j.csbj.2022.09.015.

Miller, J.H. (1972) Experiments in Molecular Genetics. Cold Spring Harbor Laboratory.

Olson, D.G. et al. (2015) ‘Identifying promoters for gene expression in *Clostridium thermocellum*’, Metabolic Engineering Communications, 2, pp. 23–29. Available at: 10.1016/j.meteno.2015.03.002.

Riley, L.A. and Guss, A.M. (2021) ‘Approaches to genetic tool development for rapid domestication of non-model microorganisms’, Biotechnology for Biofuels, 14(1), p. 30. Available at: 10.1186/s13068-020-01872-z.

Rodionov, D.A. et al. (2021) ‘Transcriptional Regulation of Plant Biomass Degradation and Carbohydrate Utilization Genes in the Extreme Thermophile *Caldicellulosiruptor bescii*’, mSystems, 6(3), p. 10.1128/msystems.01345-20. Available at: https://doi.org/10.1128/msystems.01345-20.

Straub, C.T. et al. (2020) ‘Metabolically engineered Caldicellulosiruptor bescii as a platform for producing acetone and hydrogen from lignocellulose’, Biotechnology and Bioengineering, 117(12), pp. 3799–3808. Available at: 10.1002/bit.27529.

Streett, H., Charubin, K. and Papoutsakis, E.T. (2021) ‘Anaerobic fluorescent reporters for cell identification, microbial cell biology and high-throughput screening of microbiota and genomic libraries’, Current Opinion in Biotechnology, 71, pp. 151–163. Available at: 10.1016/j.copbio.2021.07.005.

Tanwee, T.N.N. et al. (2023) ‘Metabolic engineering of Caldicellulosiruptor bescii for 2,3-butanediol production from unpretreated lignocellulosic biomass and metabolic strategies for improving yields and titers’, Applied and Environmental Microbiology, 90(1), pp. e01951–23. Available at: 10.1128/aem.01951-23.

Tjo, H. et al. (2025) ‘Maltodextrin transport in the extremely thermophilic, lignocellulose degrading bacterium *Anaerocellum bescii* (f. *Caldicellulosiruptor bescii*)’, Journal of Bacteriology, 207(5), pp. e00401–24. Available at: 10.1128/jb.00401-24.

Williams-Rhaesa, A.M. et al. (2018) ‘Engineering redox-balanced ethanol production in the cellulolytic and extremely thermophilic bacterium, *Caldicellulosiruptor bescii*’, Metabolic Engineering Communications, 7, p. e00073. Available at: 10.1016/j.mec.2018.e00073.

