## Supplementary Information for "A Straightforward and Robust Enzymatic Reporter System for Anaerobic Thermophiles"

**Table S1.** Oligonucleotide primers and ultramers used in this study.

| Primer Name | Sequence (5'→3') | Use |
|---|---|---|
| JLG021 | aactactcaccaaacctccttg | pJLG091 and pJLG093 construction |
| JLG022 | taaaggaggactataataaagggagc | pJLG091 and pJLG093 construction |
| JLG181 | gaatctcgagtcttctgacgct | Colony PCR of pSBS4 based vectors |
| JLG224 | gtgagttatacacagggctgg | Colony PCR of pSBS4 based vectors |
| JLG211_CTS480 | tcttgctctccttaaatccttgtg | qPCR-*A. Bescii gapdh* gene (Athe_1406) |
| JLG212_CTS481 | ggtgtaaatgaggagatgtacgac | qPCR-*A. Bescii gapdh* gene (Athe_1406) |
| JLG219 | gatatggatatggatgcgaaagaaac | qPCR-*Cm*βgal reporter gene |
| JLG220 | ccaaataatatggtctctgataatctgc | qPCR-*Cm*βgal reporter gene |
| B431.093.VM.F | atggatatttcttttccaaaatct | pJLG093_PmeI and pJLG161 construction |
| B431.093.VM.R | agcgtcagaagactcgagat | pJLG093_PmeI and pJLG161 construction |
| JGI.UM1 | gtcaaaaaacggtgcgcttactgttcgtgggctcatgggaatct cgagtcttctgacgctgtttaaacatggatatttcttttccaaaatc ttttagatttggatggtctcaggcaggatttcagtct | pJLG093_PmeI and pJLG161 construction |

**Table S2.** Synthesized genes and promoters used in this study.

| **Name/description** | **Sequence (overlap regions in bold underline)** |
|---|---|
| Codon optimized ***Pf*αgal** gene with overlaps for gibson assembly | **caaggaggtttggtgagtagtt**atgagagcattggtttttcatggaaacttgcagtatgcagaaattccaaaatctgaaatt<br>ccaaaagttattgaaaaagcatattttccaacaatttctgaattgattagaagagaaattccatttggattgaacattacaggat<br>attctttgtctttttgccaaaagatttgattgcattgattaaagaaggaattgaatctggattgattgaaattttgggaacatctt<br>atacacatgcaattttgccattgttgccattgtctagagttgaagcacagattaaaagagatagagaagttaaagaaaacatt<br>ttggaagtttctccagaaggattttggttgccagaattggcatatgatccaattattccagcaattttgagagataacaactat<br>gaatatttgtttgcagatggagaagcaatgttgttttctaaccatttgaactctgcaattaaaccaattaaaccattgtatccac<br>atttgattaaagcacagagaggagaaggattggtttatttgaactatttgttgggattgagagaattgaaaaaagcaattaac<br>ttggtttttgaaggaaaagttacattggaagcagttaaagaaattgaagcaattccagtttgggtttctattaacacagcagtt<br>atgttgggagcaggaagatttccattgatgaacccaaaaaaagttgcaaaatgggttaaagaaaaagatgaaattttgttgt<br>atggaacagatattgaatttttgggatatagagatattgcaggatataaaattacaatttctaacttgttggaaattattaacga<br>attggaaggagaattgggattgccaagaaaaattaaacattctgaaaaaaaattgtatttgagaacatcttcttgggcacca<br>gataaatctttgagaatttggacagaagatgaaggaaacgcaagattgaacatgttgacatcttatatggatggagaattgg<br>catttttggcagaaaactctgatgcaagaggatgggaaccattgccagaaagaagattggatgcatttaaagcaatttatac<br>acattggagatctgaaaacggaaaacatcatcaccaccaccaccat**taaaggaggactataataaagggagc** |
| Codon optimized ***Cm*βgal** gene with overlaps for gibson assembly | **caaggaggtttggtgagtagtt**atggatatttcttttccaaaatcttttagatttggatggtctcaggcaggatttcagtctga<br>aatgggaacaccaggatctgaagatccaaacacagattggtatgtttgggttcatgatccagaaaacattgcatctggatt<br>ggtttctggagatttgccagaacatggaccaggatattggggattgtatagaatgtttcatgataacgcagttaaaatggga<br>ttggatattgcaagaattaacgttgaatggtctagaatttttccaaaaccaatgccagatccaccacagggaaacgttgaag<br>ttaaaggaaacgatgttttggcagttcatgttgatgaaaacgatttgaaaagattggatgaagcagcaaaccaggaagca<br>gttagacattatagagaaatttttctgatttgaaagcaagaggaattcattttattttgaacttttatcattggccattgccattgt<br>gggttcatgatccaattagagttagaaaaggagatttgtctggaccaacaggatggttggatgttaaaacagttattaacttt<br>gcaagatttgcagcatatacagcatggaaatttgatgatttggcagatgaatattctacaatgaacgaaccaaacgttgttca<br>ttctaacggatatatgtgggttaaatctggatttccaccatcttatttgaactttgaattgtctagaagagttatggttaacttgat<br>tcaggcacatgcaagagcatatgatgcagttaaagcaatttctaaaaaaccaattggaattatttatgcaaactcttcttttac<br>accattgacagataaagatgcaaaagcagttgaattggcagaatatgattctagatggatttttttttgatgcaattattaaagg<br>agaattgatgggagttacaagagatgatttgaaaggaagattggattggattggagttaactattattctagaacagttgtta<br>aattgattggagaaaaatcttatgtttctattccaggatatggatatggatgcgaaagaaactctatttctccagatggaagac<br>catgctctgattttggatgggaattttatccagaaggattgtatgatgttattatgaaatattggtctagatatcatttgccaattt<br>atgttacagaaaacggaattgcagatgcagcagattatcagagaccatattatttggtttctcatatttatcaggtttatagag<br>caattcaggaaggagcaaacgttaaaggatatttgcattggtctttgacagataactatgaatgggcatctggattttctatg<br>agatttggattgttgcaggttgattattctacaaaaaaacagtattggagaccatctgcatatgtttatagagaaattgcaaaa<br>tctaaagcaattccagaagaattgatgcatttgaacacaattccaccaacaagatctttgagaagacatcaccaccaccac<br>cat**taaaggaggactataataaagggagc** |
| Bifurcating hydrogenase promoter (**$P_{bh}$**) with overlaps for gibson assembly | **gctcatgggaatctcgagtcttctgacgct**tccattcctcagatgcccatcatctatgggagataaatgaaagggaatttt<br>ttattgaaagtgatatactgtatacaatatttttcaattaaattctccaaaatttatacttcatttataacccgttgtatgctacaata<br>ttaacagtggttttaactccatatgttaaatttctaacaatagaagggggatgcagattt**atggatatttcttttccaaaatct<br>tttaga** |

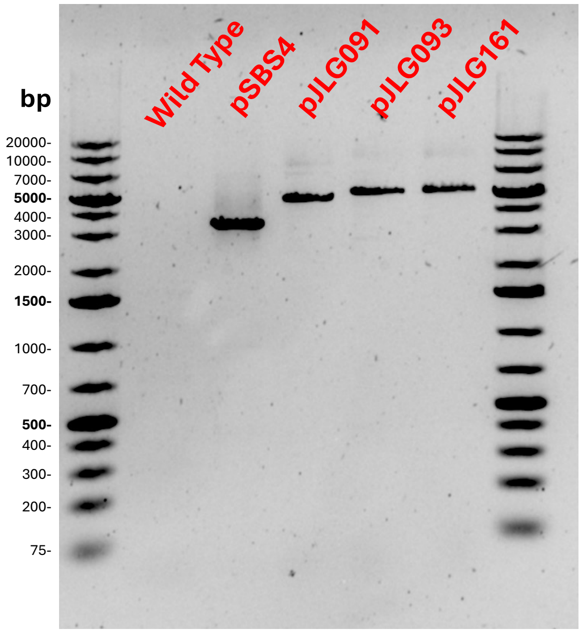


**Figure S1.** Colony PCR of strains used in this study (**Table S1; Primers JLG181 & 224**) . Wild type *A. bescii* DSM 6725 cells show no amplification as expected. Observed bands align with expected amplificon sizes of 3.3 kb for pSBS4, 4.7 kb for pJLG091, and 5.1 kb for pJLG093 and pJLG161.

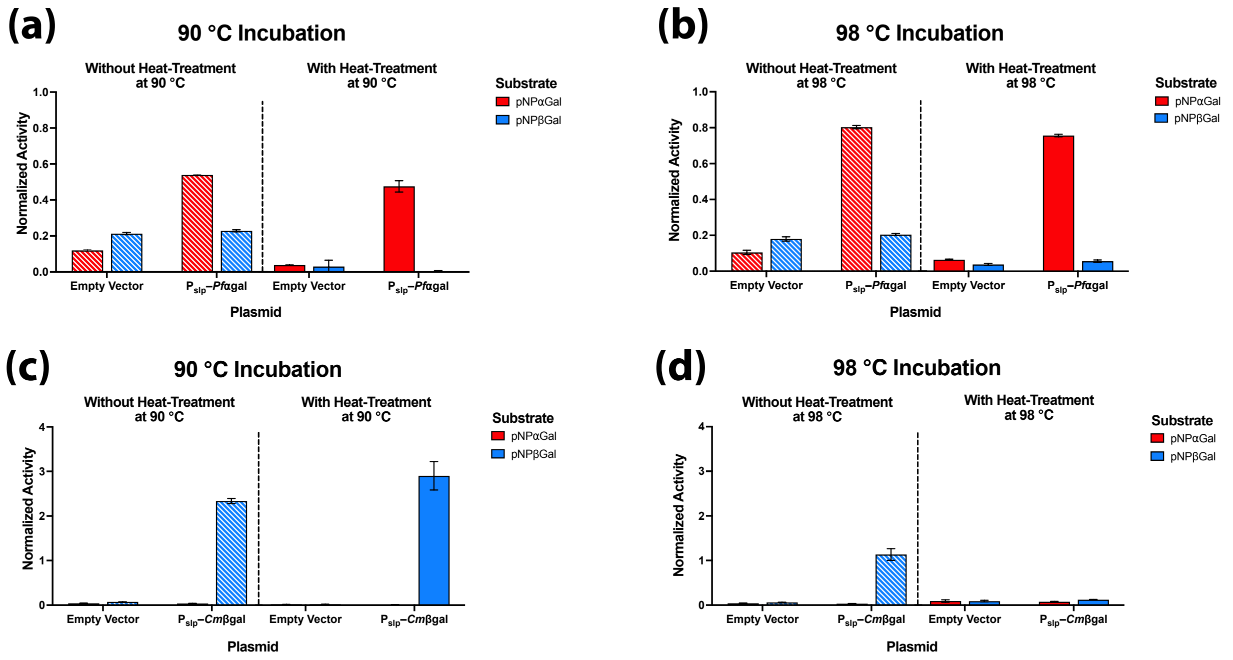


**Figure S2**. Activity of *Pf*αgal and *Cm*βgal on both pNP substrates at various reaction conditions to assess expression of these hyperthermophilic galactosidases as reporters in *A .bescii.* Here expression of both reporters is driven by $P_{slp}$. Activity of *Pf*αgal vs. the Empty Vector strain on pNPαGal and pNPβGal for: **(a)** 2 hours at 90 °C with and without 10 minutes of heat-treatment at 90 °C; **(b)** 2 hours at 98 °C with and without 10 minutes of heat-treatment at 98 °C. Activity of *Cm*βgal vs. the Empty Vector strain on pNPαGal and pNPβGal for: **(c)** 20 minutes at 90 °C with and without 10 minutes of heat-treatment at 90 °C; **(d)** 20 minutes at 98 °C with and without 10 minutes of heat-treatment at 98 °C. Error bars in all panels represent one standard deviation calculated from triplicate technical replicates at each reaction condition.
